# Information theory and the phenotypic complexity of evolutionary adaptations and innovations

**DOI:** 10.1101/070854

**Authors:** Andreas Wagner

## Abstract

Two main lines of research link information theory to evolutionary biology. The first focuses on organismal *phenotypes*, and on the information that organisms acquire about their environment. The second connects information-theoretic concepts to *genotypic* change. The genotypic and phenotypic level can be linked by experimental high-throughput genotyping and computational models of genotype-phenotype relationships. I here use a simple information-theoretic framework to compute a phenotype’s information content (its phenotypic complexity), and the information gain or change that comes with a new phenotype. I apply this framework to experimental data on DNA-binding phenotypes of multiple transcription factors. Low phenotypic complexity is associated with a biological system’s ability to discover novel phenotypes in evolution. I show that DNA duplications lower phenotypic complexity, which illustrates how information theory can help explain why gene duplications accelerate evolutionary adaptation. I also demonstrate that with the right experimental design, sequencing data can be used to infer the information gain associated with novel evolutionary adaptations, for example in laboratory evolution experiments. Information theory can help quantify the evolutionary progress embodied in the discovery of novel adaptive phenotypes.

## Introduction

The language of information permeates many different areas of biology [1-3]. In genetics, genes “encode” proteins, and RNA molecules carry “messages” from genes to protein-manufacturing ribosomes. In cell biology, “signals” are exchanged between cells. In developmental biology, organisms execute “programs” as they develop from zygote to adult [2, 4]. Information theory has also been attractive to evolutionary biologists [2, 3, 5-7]. That’s because the survivors of natural selection harbor genetically-encoded information about their environment. It is this information that renders them well-adapted to the environment.

Among the first researchers to explore the link between information and evolution was Motoo Kimura. He built on earlier work by J.B.S. Haldane to argue that adaptive evolution accumulates genetic information in proportion to the rate at which alleles are replaced by better-adapted alleles [8, 9]. Consistent with established information theoretic concepts [10], he expressed genetic information as the binary logarithm (log_2_) of the total number of possible DNA strings with the same length (in base pairs) as the human genome.

More recently, two independent lines of research have connected evolutionary biology and information theory. The first is centered on *phenotypes*, and asks how organisms enhance their fitness through phenotypes that harbor or collect information about the environment. An example of such a phenotype is the growth rate of bacteria. To regulate this growth rate, bacteria need to acquire information about available nutrients in their environment. Another is pigmentation in butterflies, and especially the dark coloration of melanic forms. Melanism helps butterflies stay warm in cold temperatures, and its development can depend on information about environmental temperature. Yet another example is the fraction of seeds that germinate in any one year in populations of desert annual plants. For optimal reproductive success, this phenotype needs to incorporate information about past rainfall patterns [5, 11-21].

A second line of research focuses on *genotypes* [3, 22-27]. Equations from classical population and quantitative genetics describe how genotypes and fitness change in evolution by natural selection, mutation, and genetic drift. These equations can also be written using information-theoretic concepts, such as Shannon’s entropy [3, 27]. This line of research shows that the distribution of a population’s allele frequencies encodes information, and that selection can increase this information [3, 25]. Unfortunately, marrying population genetics with information theory can require complex mathematics. What is more, current models to derive stationary allele frequency distributions of evolving populations require restrictive assumptions, such as that each locus harbors only two alleles, that different loci are in linkage equilibrium, and that selection and drift are weak [22].

Recent technological advances in DNA sequencing and microarrays allow us to genotype many organisms or evolving molecules. In doing so, they can also provide phenotypic information, and thus help link genotypes and phenotypes. One example is the ability of specific proteins to bind transcriptional regulators and thus regulate gene expression, which can be quantified with protein-binding microarrays [28, 29]. With technologies like these in mind, I will here use a simple information theoretic framework to quantify the informational complexity of an organismal phenotype, such as the ability to regulate a gene in a new way, or to survive on a novel nutrient.

The framework also allows me to take an information-theoretic perspective on the distinction between evolutionary adaptations and innovations [30]. While many evolutionary adaptations involve the *quantitative* fine-tuning of existing phenotypes, such as the rate at which an enzyme catalyzes a metabolic reaction, some are *qualitatively* new adaptations – innovations. Examples include an organism’s ability to thrive on previously toxic molecules, such as antibiotics, or to use new sources of carbon and energy [31, 32]. Innovations could in principle emerge through single mutations or through multiple small (and perhaps individually adaptive) mutations. A rigorous conceptual distinction between adaption and innovation is difficult [30]. I thus ask whether one can better distinguish the two concepts in information-theoretic terms.

The framework applies to all evolutionary processes, but especially to experimental evolution, where changes in genotypes and phenotypes are easily monitored. Three kinds of experimental evolution techniques are germane. The first is *in vitro* selection [33]. Here, large collections (“libraries”) of DNA or RNA genotypes are synthesized, and molecules with specific phenotypes, such as the ability to bind ATP, are selected via methods like affinity chromatography [34-37]. A variant of this approach is found in protein-binding microarrays, where a library of DNA molecules is immobilized on an array. By exposing the array to a transcriptional regulator, one can identify DNA molecules that are bound by this protein and that could mediate gene expression regulation by this protein [28, 29, 38].

A second approach is the directed experimental evolution of individual proteins or RNA molecules. Here, populations of molecules undergo repeated cycles of mutation and selection for a desired phenotype, such as the ability to cleave an antibiotic [39–48]. Directed evolution can proceed *in vitro, in vivo*, or in both. For example, one can encode the gene for an antibiotic resistance protein on a plasmid, which can then be subject to replication and selection in a host organism. Mutagenesis can be applied either *in vitro*, for example through an error-prone polymerase chain reaction (PCR) or in the host organism [44, 45, 49].

A final technique involves the experimental evolution of whole organisms, such as bacteria, algae and viruses [50-59]. As opposed to directed evolution, whose goal is often to identify molecules with specific catalytic or ligand binding activities, experimental evolution often aim to study the evolutionary process itself. It asks, for example, how organisms adapt to specific nutrients [54], stressors [52], or cyclically varying environments [54].

In all three techniques high throughput-sequencing can be applied to post-selection or post-evolution populations. The resulting sequence data can help identify the genotypes that are associated with a phenotype of interested, which is central for the approach I pursue here.

In the next section, I will first introduce the information-theoretic framework, and illustrate its use to understand evolution by DNA or gene duplication. Second, I will apply the framework to a small and tractable genotype space of transcription factor binding sites that has been characterized exhaustively through protein binding microarrays [28, 29, 38]. Third and finally, I will show how sequence data obtainable from laboratory selection or evolution experiments with today’s technology can help quantify differences in the amount of information gained by different evolutionary adaptations. In this context, I will also discuss future challenges. They regard experimental designs and statistical theory suitable to estimate information gains from sequence data.

## Results

### Concepts

A genotype space is a collection of genotypes, such as all DNA sequences of a given number of nucleotides or base pairs. Genotypes are ultimately DNA or RNA sequences, but other representations, such as amino acid sequences [60], are used for some purposes.

Genotypes are dwarfed in their diversity by phenotypes, which range from an organism’s body plan, to a metabolism’s ability to synthesize a specific spectrum of biomass molecules, and an amino acid string’s ability to fold in three dimensions and catalyze a specific chemical reaction.

Biological evolution takes place in a genotype space, where evolving populations of organisms or molecules change their genotypes through processes like mutation and recombination, and where selection allows the survival of well-adapted phenotypes. The advantage of the genotype-space perspective is that it is comprehensive. A genotype space of protein-coding DNA sequences encodes all possible phenotypes that amino acid strings can form, and thus also every conceivable protein function, be it in catalysis, transport, motility, or structural support. An evolving population that acquires any novel protein-based adaptation has discovered a genotype with a specific ability in this space. Genotype spaces are the places where all adaptations and innovations take place. They can be very large, comprising for example more than 20^100^=1.3×10^130^ proteins of merely 100 amino acids, thus making their exhaustive mapping infeasible.

The relationship between genotypes and phenotypes – the genotype-phenotype map – has been studied for multiple different kinds of genotype spaces, either exhaustively (for small spaces) or through random sampling, using both computational and experimental techniques [33, 34, 61-67]. Among the important insights from such analyses is that, first, (astronomically) many genotypes usually form the same phenotype. Second, these genotypes are organized into one or more networks in genotype space, where network members can be reached from each other through multiple small genotypic changes that leave the phenotype intact. Third, the genotype networks of different phenotypes are interwoven in complex ways [64, 67, 68]. Fourth, some phenotypes have larger associated genotype networks than others, which also extend farther through genotype space. With possible exceptions [69, 70], populations evolving on a large genotype network have a greater potential to stumble upon new and beneficial phenotypes, because such populations can explore a larger proportion of genotype space [67, 71-73].

My starting point is the observation that for any observed phenotype *P*, such as a protein’s ability to bind or react with a specific molecule, there are usually multiple genotypes *G_P_* encoding this phenotype. To simplify the exposition, I focus on qualitative (binding or not) rather than quantitative phenotypes (e.g., binding with a specific affinity). That is, I assume that all genotypes with a particular phenotype are equivalent, and when sampling from a specific set of such genotypes, the probability *p(g)* of drawing any one genotype is the same for all genotypes *g.* This assumption can be relaxed.

The amount of information in a single genotype *g* drawn at random from a genotype space *G* is given by the Shannon entropy of a random variable whose possible values are all genotypes *g* in *G*, and that assumes these values with equal probability 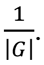 I denote this information content as *H(G)*, and it computes as 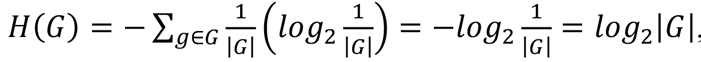 where |*G*| denotes the number of genotypes in the set *G.* Analogously, the information content for a member *g* of a subset *G_P_* of genotypes with a specific phenotype *P* (Figure 1a), computes as

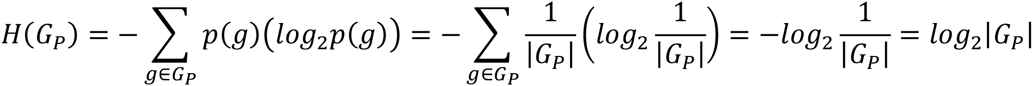

These observations give rise to the following definition.

**Figure 1.**
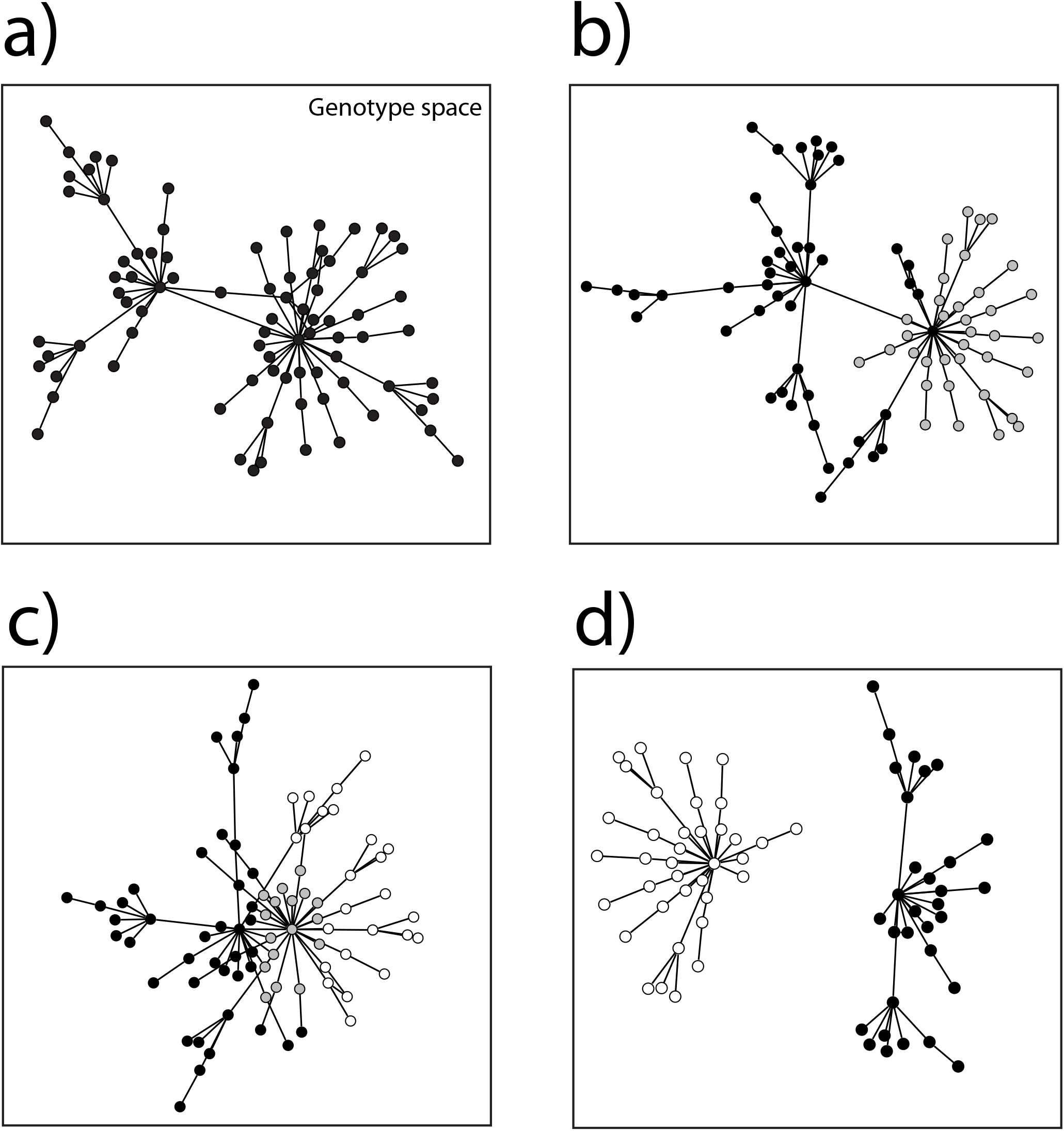
Genotype sets in genotype space can have various topological relationships. Large rectangles symbolize genotype space, circles correspond to genotypes, and straight lines connect genotypes that are (1-mutant) neighbors, i.e., they differ by a small genetic change such as a single nucleotide change. Each set is shown as a network, because genotype sets usually form networks in genotype space. **a)** A hypothetical set of genotypes with the same phenotype. The set is shown as a single genotype network, but I note that it could consist of multiple disconnected networks. **b)** Two sets of genotypes, each corresponding to one phenotype, an old phenotype (black *and* grey circles), and a new phenotype (grey circles only). The genotypes with the new phenotype form a subset of genotypes with the old phenotype. **c)** Sets of genotypes with two phenotypes, an old phenotype (black circles), a new phenotype (white circles), and genotypes with both the old and new phenotype (grey circles). Unlike in b), the genotype set of the new phenotype is not a subset of the genotype set with the old phenotype. **d)** The sets of genotypes for different phenotypes can be non-overlapping.

#### Definition 1

The amount of information that is associated with phenotype *P* is given by

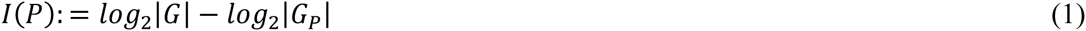

Following analogous uses in DNA sequence analysis [74], I will also refer to this quantity as a phenotype’s (informational) *complexity.* The greater this complexity, the more information is required to encode the phenotype. When applied to aptamers and ribozymes, this quantity has also been called functional information [75]. Because function is an ambiguous word, and because not all phenotypes may have a function, I prefer to use the more general notion of phenotypic information.

Some empirical data on phenotypic complexity is available for macromolecules. For example, in vitro selection experiments identifying ATP-binding proteins from a random protein library with 80 amino acids have estimated that a fraction *f_P_*=10^-11^ or 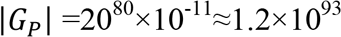 proteins of this length can bind ATP [34]. Individual proteins in this set thus harbor *log_2_*|1.2 × 10^93^| = 309.2 bits of information. The amount of information associated with the ATP-binding phenotype is 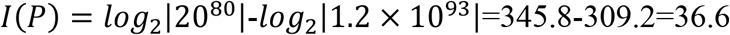 bits. To compare data from genotype spaces of different size (e.g., different lengths of proteins), it can be useful to consider the number of bits per monomer, which in this example is 36.6/80=0.46, or 10.6 percent of the maximally possible value of 345.8/80=4.32 bits per amino acid. (This fractional information can also be computed as 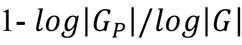, regardless of the base of the logarithm.)

Unlike with in vitro selection experiments, laboratory evolution experiments often do not start from random collections of genotypes, but from genotypes that already have a specific phenotype *P_Old_* and acquire a novel phenotype *P_New_* (Figure 1b). For example, in a directed evolution experiment, TEM-1 β-lactamase molecules that convey resistance to ampicillin may acquire the ability to also cleave the antibiotic cefotaxime. Denote as *G_Old_* the subset of genotypes with the old phenotype, and as *G_New_* the subset of genotypes with the new phenotype. To obtain the information gain associated with the new ability, it is useful to consider the relative entropy (or Kullback-Leibler distance, [76]) of two random variables *O* and *N* that are defined on the genotype space G. For all genotypes *g∈ G_Old_*, the probability that the random variable *O* assumes a value of *g* is equal to 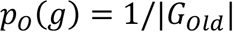 and for all genotypes 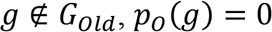. Likewise, for a genotype with the new phenotype *g∈ G_New_*, the probability that the random variable *N* assumes a value of *g* is equal to 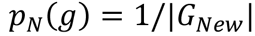 and for 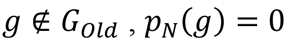. In other words, *O* and *N* are indicator variables that indicate the membership of a genotype in the sets *G_Old_* and *G_New_*. The relative entropy of the two random variables is then given by

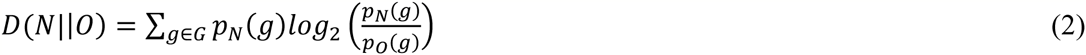

The relative entropy is always greater than zero, except when 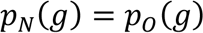 for all genotypes, which requires that *G_New_ = G_Old_*. In that case, *D(N||O)* = 0. Despite the suggestive name Kullback-Leibler *distance, D* is not a true distance, because it is not symmetric [76]. Specifically, for the random variables considered here, and whenever *G_New_* is a proper subset of G_Old_, *D(O||N)* = ∞, because there will be some genotypes that are in G_Old_ but not in *G_New_*, and for these genotypes 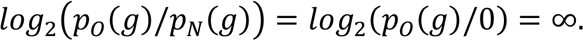

To simplify equation (2), I take advantage of the conventions 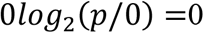 and 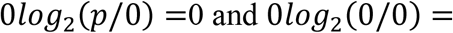=0[76 p.19] to obtain

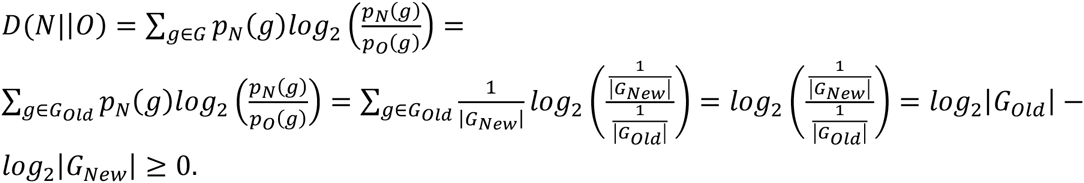

From these observations, one can define the information gain associated with the acquisition of a new phenotype *P_New_* starting from some phenotype *P_Old_*, where *G_New_⊆G_Old_* as

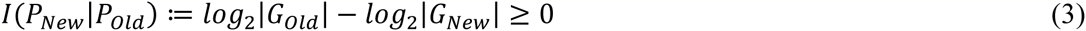

In other words, this information gain is equivalent to the difference in information content between members of the old and new sets of genotypes. The information content of a phenotype from definition 1 (equation 1) can also be viewed as a Kullback-Leibler distance. If one expresses the number of genotypes in *G_New_* as a fraction *f_New_* ≤ 1 of those in *G_Old_*,i. e., *|G_New_| = f_New_|G_Old_|*, then the relative information gain becomes 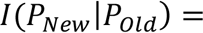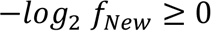.

The situation is different when *G_New_* is not a proper subset of *G_Old_* (Figure 1c). In the hypothetical β-lactamase example, this scenario might correspond to enzymes that have the new phenotypic ability to inactivate cephotaxime, but that may or may not have retained the old phenotype to inactivate ampicillin. In this case, the Kullback-Leibler distance is undefined, because there will be some genotypes *g* ∈ *G_New_* that are not also in *G_Old_*, and for these genotypes the summands in equation (2) take the form 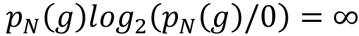. However, one can still calculate the difference *I(P_New_|P_Old_)* from equation (3), except that this difference need not be greater than zero. It can be smaller than zero if, for example, more genotypes are able to cleave cefotaxime than ampicillin. In other words, the acquisition of a new phenotype may incur a net information or complexity loss, if loss of an old phenotype is permitted. Overall, we arrive at

#### Definition 2

The information or complexity *change* associated with the acquisition of a new phenotype *P_New_* starting from some phenotype *P_Old_*, is given by

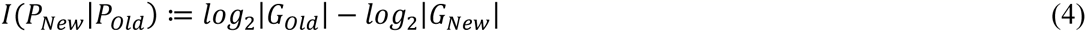

The above definitions can be extended to scenarios where genotypes in a genotype space *G* (or a set of genotypes *G_P_)* are not equivalent, by replacing the above uniform probabilities of sampling a genotype from a given set by non-uniform probabilities. This can occur for at least two reasons. First, the processes generating individual genotypes may not generate them with equal probability. For example, oligonucleotide synthesis or mutagenesis can cause non-uniform nucleotide compositions in random DNA libraries used for *in vitro* selection experiments [77]. Mutations in living organisms show such biases as well, for example in the well-known higher frequency of transition (A↔T, G↔C) to transversion mutations [78]. The second reason applies when one considers quantitative phenotypes, such that not all genotypes in a set *G_P_* may express the phenotype *P* to an equal degree. For example, different proteins may bind a ligand like ATP with different affinity. As a result, some proteins are preferentially selected during experimental evolution or selection for this affinity. The consequence in both cases is that the probability *p(g)* of observing a specific genotype is no longer uniform. What is more, this probability then depends not only on the phenotype itself, but on details of experimental design, such as mutation rates, population sizes, and selection strength. I will restrict myself to the uniform case corresponding to unbiased sets of genotypes and qualitative phenotypes.

### Application to DNA or gene duplication

Because of its simplicity, this framework can help shed light on various questions associated with the origin of evolutionary adaptations and innovations. To give but one example, consider the duplication of DNA and its role in evolutionary adaptation. It has long been thought that such duplications, and in particular duplications that include entire genes, can accelerate the origin of novel phenotypes [79-83]. An evolutionary advantage associated with duplication may exist, because an evolving duplicate stretch of DNA can explore a larger genotype space (or a larger proportion of such a space) than either could alone, while preserving its pre-duplication phenotype. An information theoretic framework may help quantify this advantage.

Consider some phenotype, such as a regulatory region’s ability to bind a specific transcription factor, or a protein’s ability to catalyze a specific chemical reaction, and the set of genotypes *G_P_* associated with this phenotype, which occupies a fraction *f_P_* of genotype space. When the DNA encoding this phenotype becomes duplicated, both copies can undergo DNA mutation independently. Thus, they evolve in a larger genotype space, which is the Cartesian product space of *G (G × G*) and comprises many more (|G|^2^) genotypes. If each genotype is *L* nucleotides long, the duplicated genotype contains twice as much information (log_2_|G|^2^=2log^2^|G|, but the same amount of information per nucleotide 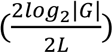, as before the duplication 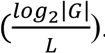 How many genotypes with phenotype *P* are there in the new genotype space? If it was necessary that both copies have this phenotype, their number would simply be given by |G_P_|^2^. However, only one of the copies may need to have the phenotype. In this case, the set of genotypes becomes much larger, because it is comprised of three subsets. The first of them comprises all genotypes where both copies have the phenotype. The second is the set of genotypes where the first of the two copies has the phenotype. It is given by *G_P_* × *(G|G_P_)*, where “\” denotes the set-theoretic difference *(G* without *G_P_*), and has size |*G_p_*|(1 — *f_P_)|G|*. The third is the set of genotypes where the second of the two copies has the phenotype, *(G\G_P_) × G_P_*, and it has the same size. Adding the members of the three sets yields the following number of genotypes

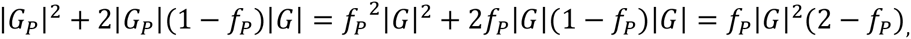

and calculating the ratio of this number to the number of genotypes in which both copies have phenotype *P* yields

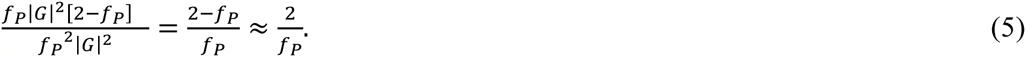

The right-most approximation holds for the typical case where *G_P_* occupies a tiny fraction of genotype space, such that *f_P_* is orders of magnitude smaller than unity. In terms of the ATP-binding protein above, where |*G*| =20^80^ and *f_P_*=10^-11^, the admissible genotype set after a gene duplication is a factor 2/10^-11^≈2×10^11^ larger than if both genotypes need to encode an ATP-binding phenotype. This indicates how great the advantage of gene duplication can be in terms of the number of new genotypes that can be explored. Because this number scales as 1/*f_P_*, this advantage will increase as *f_P_* decreases. In other words, it will be greatest for phenotypes that are formed by only a small set of genotypes.

The difference in informational complexity before and after duplication computes as

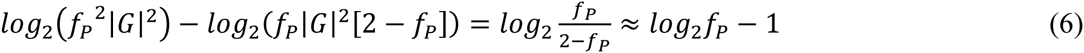

as long as *f_P_* is small. An equivalent expression is obtained if one compares the respective sets in the pre-and post-duplication genotype spaces. Because the dimension of genotype space is changed as a result of the duplication, an appropriate comparison requires the per-residue information content. Specifically,

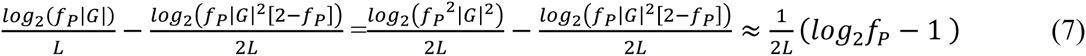

Note that in both comparisons, the difference in information content is negative, because the set of post-duplication genotypes where at least one genotype has the required phenotype occupies a larger fraction of genotype space, and thus contains less information – its informational complexity is lower. This is important, because exploring a larger sets of genotypes (containing less information) facilitates the exploration of novel phenotypes that can be found near this set [67, 71-73]. In terms of the ATP-binding protein above, where *L*=80, the information content of each sequence changes by 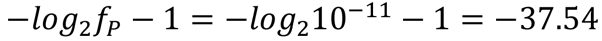 bits or by - 37.54/(2×80)≈0.23 bits per residue.

### Transcription factor binding phenotypes

I next illustrate how these concepts can be applied to the genotype space of all 4^8^ = 65,536 DNA sequences of length eight nucleotides. Protein binding microarrays that harbor all such sequences have been used extensively to measure the binding of individual transcription factors to such sequences. Numerous evolutionary adaptations and innovations have been associated with the origin of transcription factor binding sites on DNA [84-87]. They range from changes that affect the virulence of microbial pathogens [85], to adaptive pigmentation changes in multicellular organisms [86], to more profound changes in body plan, such as the evolution of two-winged from four-winged insects, and the transformation of one wing pair into a balancing organ [84, 86].

To estimate the amount of information gained through a novel adaptation whose genetic basis is a new transcription factor binding site (and thus usually a novel pattern of gene regulation), I use previously published protein binding microarray data from 187 mouse transcription factors on the binding of each factor to all 4^8^ sequences in sequence space. My analysis is qualitative and focuses on all sequences that are bound with high affinity (see Methods). The emergence of a new binding site can either take place *de novo* or from a pre-existing binding site. For *de novo* emergence, the relevant phenotypic complexity is the information content of the binding site (definition 1), where the phenotype in question is the ability to bind a specific transcription factor. Figure 2a shows a histogram depicting this phenotypic complexity for all 187 transcription factors, and in the inset, the fractional volume of genotype space that each factor can bind, which ranges from 0.0016 to 0.045. The phenotypic complexity of transcription factor binding ranges widely, from 4.48 to 9.31 bits (median: 5.72) bits, and is proportional to the specificity of a transcription factor. Highly specific transcription factors bind fewer sites, and the ability to bind them requires more information. Most transcription factors in the data set have binding sites that are eight nucleotides long or shorter [28, 29, 88]. The shorter a binding site is, the more eight-mers a factor will bind, and the smaller the complexity of the binding phenotype.

**Figure 2.**
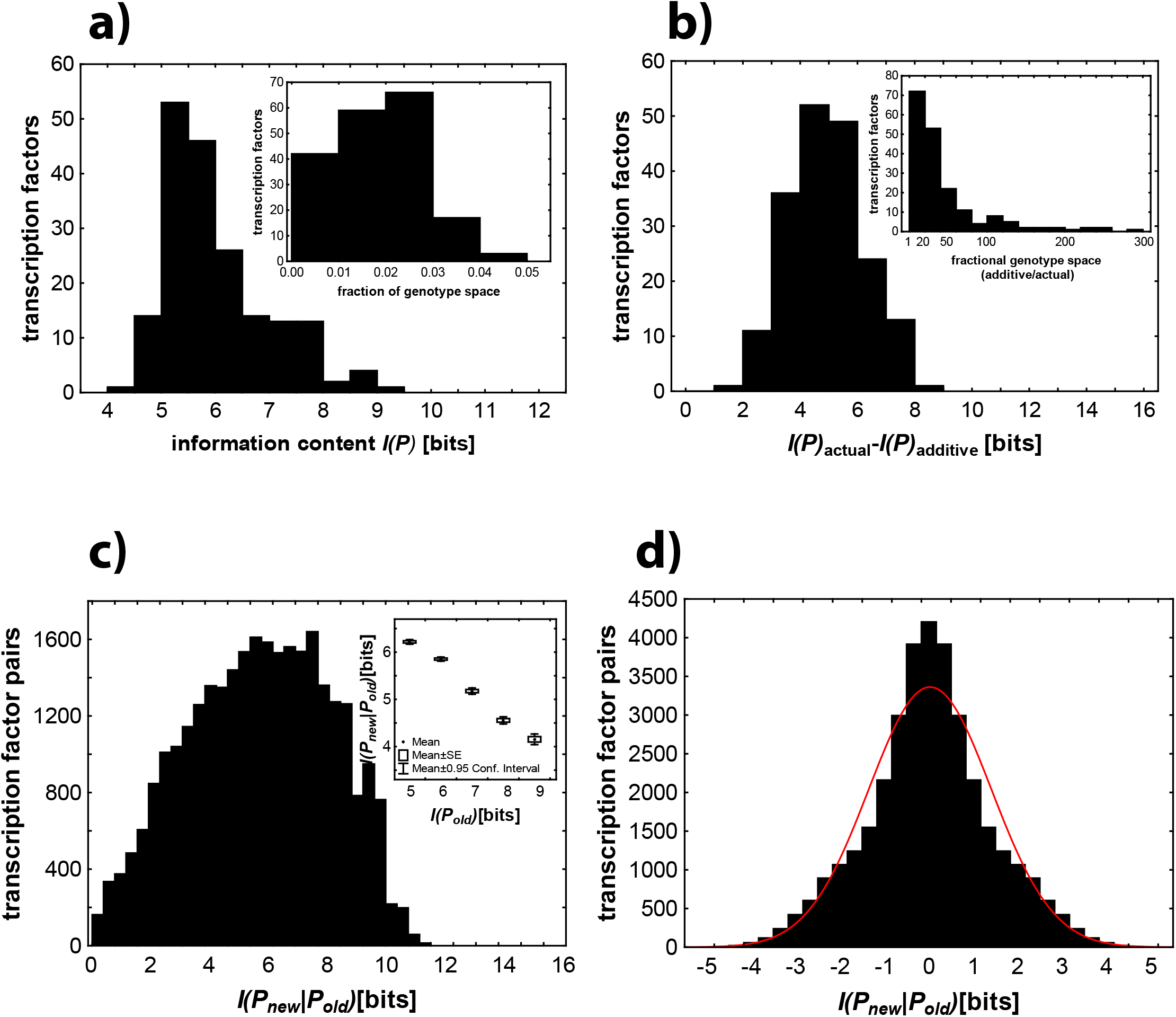
Information content and change associated with the acquisition of transcription factor binding. Data in all panels are based on experimentally measured binding of 187 mouse transcription factors to all possible DNA binding sites of length eight [28, 29, 88]. **a)** Histogram of the information content of the DNA binding phenotype of each transcription factor (definition 1 and equation 1). The inset shows the distribution of the fraction of genotype space occupied by all binding sites bound by each factor. **b)** Histogram of the difference between the actual information content of a phenotype and that estimated under the assumption that individual binding site positions contribute additively to the information content. Note that this difference is always positive, i.e., the additivity assumption underestimates the information content by estimating more sites to be bound than are actually bound. The inset shows the distribution of the ratio between the actual fraction of genotype space occupied by transcription factor’s binding sites and the estimated fraction based on the additivity assumption. This ratio is always greater than one. **c)** The gain in information content associated with acquisition of a new DNA binding phenotype, when an old phenotype is simultaneously preserved (equation 3). The inset shows this gain in information content (vertical axis) as a function of the information content of the old phenotype (horizontal axis). Circles correspond to means, boxes to standard errors, and whiskers indicate 95 percent confidence intervals. Data in c) are based on all 29290 pairs of transcription factors whose sets of binding sites overlap. **d)** The change in information content associated with acquisition of a new DNA binding phenotype when the old phenotype need not be simultaneously preserved (equation 4). The red line indicates the fit to a Gaussian distribution. Data in d) are based on all 187^2^ pairs of transcription factors in the data set.

These numbers can also help illustrate the advantage that duplication of a regulatory region containing a binding site can provide, if two sites can evolve separately after duplication, and if only one site needs to preserve its DNA binding ability. According to equation 6 and the data in Figure 2a, duplication changes the information content by -5.48 to -10.31 bits (-0.34 to -0.64 per post-duplication bit, equation 7), depending on the transcription factor. This is a substantial reduction in complexity, considering that the (post-duplication) genotype space encodes only a maximum of 32 bits of information. These numbers translate into between 44.4 and 1250 times more genotypes that an evolving population of duplicated binding sites can explore without losing its DNA binding ability (equation 5). In reality, the advantage of duplication will usually be much greater, because individual binding sites are parts of larger regulatory regions, and much more DNA than an individual binding site may be affected by a duplication.

The information content of a binding site is usually estimated from a position weight matrix, which summarizes binding information from multiple bound DNA motifs to compute a binding affinity score for any one DNA sequence, under the assumption that individual nucleotides contribute additively to binding [89]. The additivity assumption will underestimate information content, i.e., it will overestimate the number of sequences bound, and thus underestimate the information gain. The reason is that some nucleotides at two or more positions of a binding site may bind strongly individually, but not in combination, and the additivity assumption cannot capture such non-additive (epistatic) effects. Figure 2b displays a distribution of the number of bits by which the additivity assumption overestimates the information gain. It ranges from 1.56 bits for transcription factor Tbp (the TATA-binding protein) to 8.22 bits for Usf1 (upstream transcription factor 1). In terms of the number of sequences bound, the additivity assumption overestimates this number by a factor that lies between 2.9 and 300, depending on the transcription factor (Figure 2b, inset). Thus, even though transcription factor binding generally involves only modest amounts of epistasis [88], this modest amount can lead to a substantial underestimation of information content. Whenever comprehensive binding information is available, it is preferably to estimate information gain from them, without computing position weight matrices.

When binding of a new transcription factor originates at a site bound by an old factor, such that binding of the old factor must be preserved (for example, because the old factor directs gene regulation in a different tissue), then phenotypic complexity increases (equation 3 and 4, definition 2). Figure 2c shows the distribution of relative entropies for all pairs of mouse transcription factors whose genotype sets of binding sites overlap. The smallest information gain (0.04 bits) occurs when a binding site for factor Myb originates from one for factor Mybl1. The reason is that these factors, which are encoded by members of the same gene family, bind to a similar number of sites (1969 and 1775, respectively), and that most of these sites are identical: 97.2% of sites bound by Mybl1 are also bound by Myb. The largest information gain (11.5 bits) occurs when binding sites for transcription factor Mnt (cisbp id: T015083) emerge from those for Sp110 (cisbp id: T139995). Sp110 binds to 2933 sites, whereas Mnt binds to a very small set of 176 sites. Moreover, there is only a single site that allows binding of both factors and that preserves the old phenotype while permitting the new one. The inset of Figure 2c shows that the information gain is generally lower if the old site had high information content (Spearman’s r=- 0.22; P<10^-17^; n=29290), illustrating that the information that can be gained with a new adaptation may depend on evolutionary history, i.e., on old adaptations and their information content.

If binding to an old transcriptional regulator need not be preserved, then a new adaptation may incur not just a gain but a loss of information. This can occur if the new regulator can bind to more sequences than the old regulator. Figure 2d shows a histogram of the amount of information change in this scenario. Its distribution is symmetric, because for every value of information change X that occurs when binding by some new regulator Y is gained and binding by an old regulator Z is lost, there is an opposite value −X when binding by Z is gained and binding by Y is lost. The values in this distribution range from a maximal information loss or gain of 4.83 bits in the transition between a binding site for transcription factor Usf1 (cisbp id: T015121 [29]; with 103 binding sites) to one for Sp110 (id: T139995; 2933 binding sites), to a minimal information change of zero bits for several factor pairs, including Hbp1 and Rfx4, both of which bind the same number of 1320 sites. The distribution of information change is leptokurtic, that is, it shows an excess of small information changes relative to a normal distribution (Figure 2d). In computing the distribution of information change, I disregarded whether a set of binding sites *G_New_* is accessible from *G_Old_* via single point mutations that do not lead outside either set (Figure 1d). In the small genotype space I consider here, the vast majority of genotype set pairs are accessible from each other. Specifically, for 99.998 percent of unique transcription factor pairs (all but 80 of 34782 pairs), the sets of binding sites either overlap, or their minimal distance in genotype space is one, that is, there exists at least one single point mutation that leads directly from *G_Old_* to *G_New_*. For the remaining 0.002 percent of site pairs, this minimal distance is two. No larger minimal distances exist in this data set.

### Inferring information content from sequence data

The DNA binding phenotypes I discussed exist in small genotype spaces, but that is not the case for data from most evolution or *in vitro* selection experiments, which explore genotype spaces of astronomical size. What is more, myriad genotypes usually encode the same phenotype, but one can only identify as many as DNA sequencing technology permits (currently thousands for microbial genomes to millions for individual genes). Thus, it will often be impossible to infer the information content of any one phenotype (definition 1). What is more, sequencing technology does not simply enumerate genotypes but *samples* them from an evolving population, which introduces further complications. However, even with such complications, it may be possible to estimate the information *gain* or *change* associated with a novel phenotype (definition 2), because this change is usually much smaller than the absolute information content. I will next highlight how this could in principle be done. In doing so, I make simplifying assumptions whose relaxation will require future work. For now, my main point is to show that quantifying phenotypic information change is within reach of current technologies.

Consider two populations of which one is well-adapted to some ancestral environment (with phenotype *P_Old_)*, and another one is adapted to a new environment, such as one that harbors a novel nutrient, an antibiotic, or another stressor, and thus requires an altered phenotype *P_New_*. I assume that both populations comprise asexually reproducing haploid individuals, that they are in mutation-selection-drift balance subject to Wright-Fisher dynamics [78], and that they have equal effective sizes *N_e_* and mutation rates μ. I also assume that both phenotypes *P_Old_* and *P_New_*are subject to strong truncation selection, that is, mutations that disrupt each phenotype are lethal. The two phenotypes may differ in their numbers of associated genotypes *G_Old_* and *G_New_*, and thus also in their information content. The task is to estimate this difference (*log_2_ | G_Old_ | — log_2_ |G_New_*) from two samples of *n* genotypes (DNA sequences), one from each of the populations. Because genotypes with the same phenotype usually form networks that extent far through sequence space, where any one genotype can suffer mutations that disrupt the phenotype [90], I model this difference as a difference in the average rate of strongly deleterious mutations across all genotypes, or equivalently, in the average rate of neutral mutations. That is, if *l_Old_* and *l_New_* denote the average proportion of all strongly deleterious (lethal) mutations in the two populations, then the average neutral mutation rate becomes 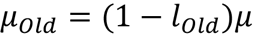 and µ_2_=(1-l_2_)µ. Assuming further that mutations in all viable sequences are equally likely to be strongly deleterious, one obtains the relationships 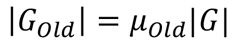 and 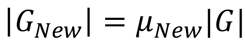, where |G| is the total size of genotype space. It is then easy to see that

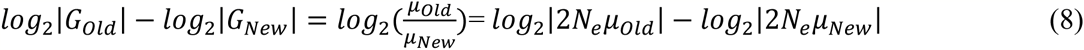

If phenotype *P_New_* harbors more information than *P_Old_* (i.e., if |*G_Old_*| > |*G_New_*|), then 2*N_eµold_*>2*N_eµNew_*. Thus, estimating the difference in phenotypic information content requires estimating the quantities 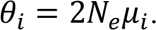 These quantities are of broad importance in population genetics because they predict a population’s amount of neutral polymorphisms [78, 91]. If phenotype *P_New_* harbors more information than 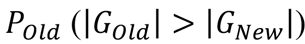 then 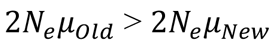 and one would expect population 2 to harbor more alleles encoding this phenotype than population 1. A maximum likelihood estimator of θ_i_ is the number of *different* genotypes *k_i_* in a random sample of *n* genotypes sequenced from the populations [92]. What is more, this number has a complex but known sampling distribution, which can be used to ask how large the difference in information content between two phenotypes must be, so that one can detect it from a sample of *n* sequences (see Methods). For example, for the null-hypothesis that phenotype *P_New_* harbors more information than *P_Old_*, one can compute the probability *p* of falsely rejecting this null-hypothesis.

(Figure 3 shows the minimally distinguishable information difference (see legend), for multiple values of the number *n* of sampled sequences, and for multiple values of 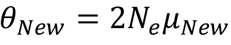. To create this plot, I chose multiple values of *n* and *θ_New_*, and for each of them I determined the smallest value of *θ_Old_* such that p<0.05. White regions in the plot indicates that for a given sequence coverage *n* and *θ_New_*, the information content of the two phenotypes is indistinguishable. In a region of the plot where the test can discriminate at least *x* bits, p<0.05 for all values of *θ_Old_*, such that 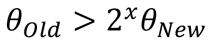.

**Figure 3.**
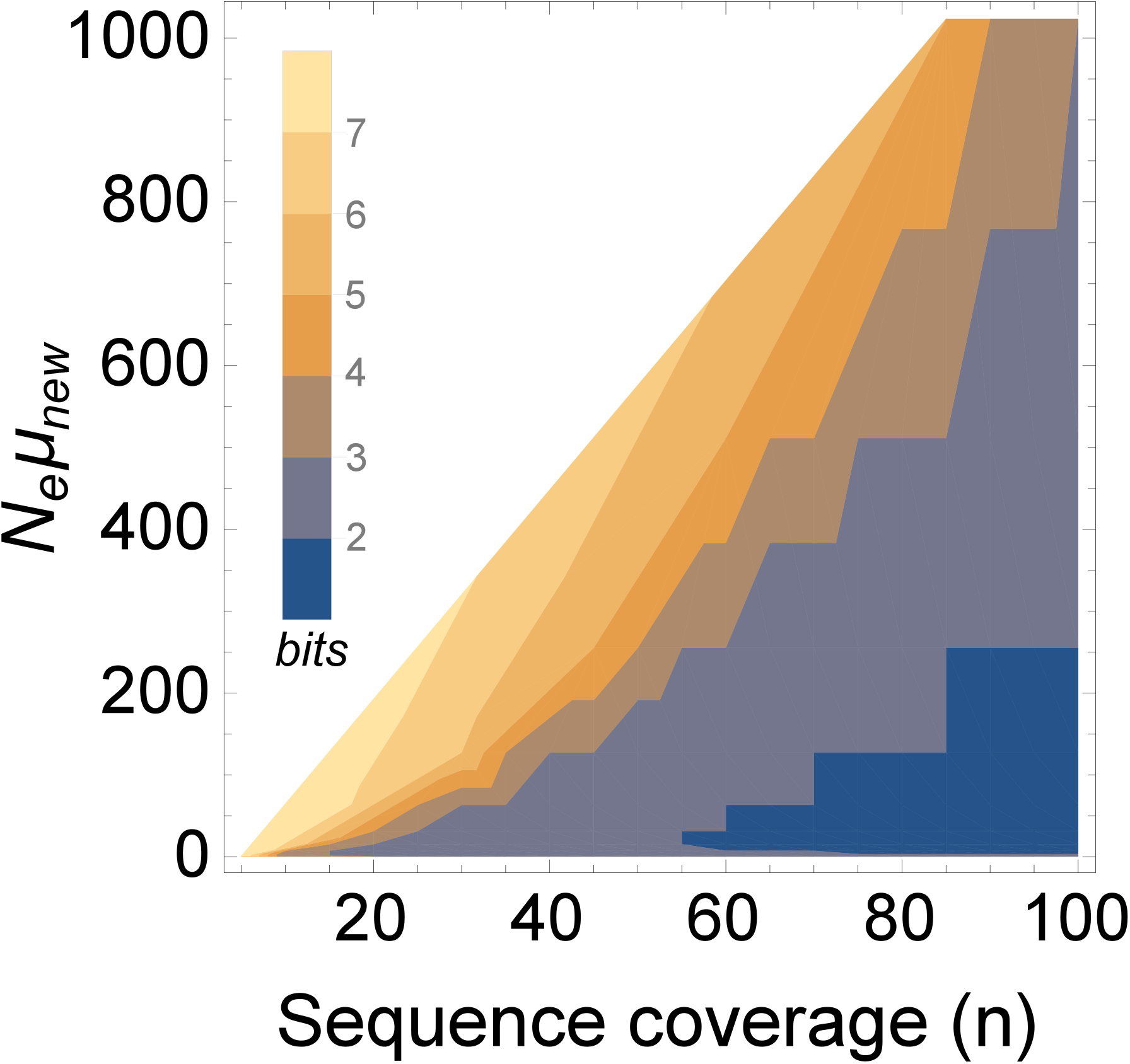
High-throughput sequencing can help distinguish even modest differences in phenotypic information content. Minimally distinguishable information content of two phenotypes (in bits), color-coded as indicated in the legend, for a given sequence coverage *n* (horizontal axis), and a given value of *θ_New_ = 2N_e_μ*_New_. To create this plot, I chose multiple values of *n* and *θ*_New_, and determined the minimal value of θ_Old_, 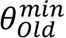 such that p=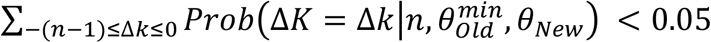 (see Methods) for each of these values. The minimally detectable information difference is then given by 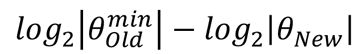.

In this analysis, I did not explore populations with *θ_Old_,θ_New_* < 1. The reason is that such populations are monomorphic most of the time [91], which implies that even when sequencing multiple genotypes, one would find mostly the same alleles. At the other extreme are values of *θ_New_ » n* (and thus also *#x03B8;_Old_ » n)*. (Figure 3 shows that in such populations, one cannot discriminate the information content of two phenotypes. The reason is that both populations are so highly polymorphic that within either population all *n* sampled sequences are likely to be different from each other. This consideration shows that sequence coverage *n* must be high relative to *θ_New_* (lower right corner of (Figure 3) for best discrimination between the information content of two phenotypes. In sum, in order to discriminate between the information content of two phenotypes, sequence coverage must be sufficiently high that 1« *_Old_,θ_New_ « n*. Again, these considerations are based on simplifying assumptions, such as strong selection, which need to be relaxed to develop a rigorous sampling theory. However, they indicate that sequence data can in principle be used to estimate differences in phenotypic information content.

## Discussion

I used a simple information-theoretic framework to quantify, first, the amount of information encoded in a phenotype and, second, the amount of information change entailed by the origin of a new phenotype. The two measures differ formally only in that information content is expressed relative to all genotypes in a genotype space, and information change is expressed relative to all genotypes with a reference phenotype. I emphasize information change and not just content for two reasons. First, in most evolution experiments, a given protein, ribozyme, or organism starts with some phenotype, and evolves a new phenotype. Second, only in exceptional small genotype spaces like those of transcription factor binding sites would it be possible to quantify phenotypic information content. In contrast, the amount of information change associated with a new adaptation may be smaller and can thus be more easily quantified experimentally. I refer to information-rich phenotypes as complex phenotypes.

It is tempting to ask about the meaning of the information embodied in a phenotype. One could argue, for example that regulatory DNA regions embody “knowledge” about the shape of the proteins that bind there. However, information theory does not require or even consider semantic aspects of information. I will thus not consider them here.

I applied the above framework to experimentally measured DNA binding affinities of 187 mouse transcription factors. Here, the phenotype is a property of a DNA molecule, i.e., its ability to bind a specific transcription factor. An advantage of this data sets is that it allows an exhaustive analysis of a tractable genotype space (6.5×10^4^ DNA sequences). In larger genotype spaces where such exhaustive exploration is not feasible, one can estimate information content under the assumption that individual nucleotides interact additively to produce a given phenotype [93]. However, this assumption generally underestimates phenotypic information content. The size of this error can be substantial, for example, up to eight out of 16 maximally possible bits for transcription factor binding sites (Figure 2b). One could correct for non-additive interactions among pairs of nucleotides [93-95], but such corrections become infeasible for the many possible higher-order interactions.

I restricted myself to qualitative or threshold phenotypes (binding/non-binding, viability/non-viability), which have proven useful in past experimental estimates of the fraction of genotypes with a particular phenotype. For example, *in vitro* selection experiments have been used to estimate that in a genotype space of RNA molecules that are 100 nucleotides long, one in 10^10^ molecules can bind to aromatic dye molecules [33], which implies a phenotypic information content of 33.2 bits. Likewise, approximately one in 10^13^ RNA molecules of length 220 encode RNA ligases, which leads to a phenotypic information content of 43.2 bits [33].

The framework can be extended to quantitative phenotypes. This requires that one replaces expressions like *log_2_|G_p_|*, which reflects equiprobable occurrence of each genotype with phenotype *P* 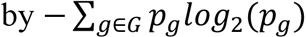 where *p*_g_reflects the probability that a specific genotype *g* occurs in a sample or population. The probability *p_g_* reflects the extent to which a phenotype is expressed by a specific genotype, e.g., the strength of binding to a transcription factor embodied in a DNA sequence. Limited empirical data is also available about the informational complexity of quantitative phenotypes. For example, it has been estimated that a 10-fold increase in an RNA aptamer’s binding affinity to guanosine triphosphate (GTP) requires 10 additional bits of information, and a 10-fold increase in an RNA ligase ribozyme’s catalytic rate requires 11 additional bits of information [75, 95].

A focus on qualitative phenotypes is a simplification, but it also facilitates an important distinction. It is the distinction between the information content of a phenotype itself, which depends only on the number of genotypes harboring this phenotype, and that of a population whose members have this phenotype [93, 94]. The latter depends not just on phenotypic information content, but also on many parameters affecting a population’s dynamics, such as (effective) population size *N_e_* and mutation rate μ. For example, if *N_e_μ«* 1, then all members of a population have identical genotypes most of the time [91]. If one were to take the one genotype sampled from such a population to calculate a phenotype’s information content, one would arrive at the maximally possible value *log_2_|G|*. However, this would be highly misleading, because there myriad genotypes might encode this phenotype, even though the population harbors only one of them. Likewise, during adaptive evolution, when a mutation has created a genotype with a superior phenotype, this genotype may sweep through the population, driving all other genotypes extinct, until it accumulates mutations and diversifies. Thus, transiently, a population’s information content can rise dramatically before reaching a mutation-selection equilibrium [93, 94]. Population sampling during the transient period would lead to misleading estimates of phenotypic information content.

The genotypes associated with a new phenotype may be difficult to access by an evolving population. Such low accessibility can have two, not mutually exclusive reasons. First, the phenotype may have high information content and thus a small number of encoding genotypes, which are difficult to find through random exploration of this space. Second, there may be many such genotypes, but a population may be far removed from them in genotype space. Multiple mutations may be required to reach them, or they may be unreachable, if some mutational intermediates are inviable. In other words, historical contingencies, for example past environments that a population encountered and adapted to, can affect the accessibility of specific phenotypes. An advantage of the information-theoretic framework is that it eliminates such historical factors from consideration, and focuses on intrinsic properties of phenotypes. A limitation is that historical factors can influence the estimation of phenotypic information content from sequence data. I note that in the data sets I analyzed here, phenotypic accessibility is not a major problem. For all pairs of transcription factors I analyzed, evolutionary transitions between binding sites for one factor to those of the other require can be achieved with single nucleotide changes. However, in larger genotype spaces, phenotypic accessibility may be more constrained [96].

I argued that one may be able to distinguish even modest phenotypic information changes in evolving populations with existing sequencing technology and the right experimental design. This argument rests on simplifying assumptions. One of them is that such populations should be in mutation-selection balance. While such balance is in practice only achieved asymptotically, it is approached exponentially with decay parameter 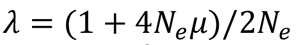 [91, p 204]. For an evolving *E.coli* population of 10^4^-10^7^ individuals with μ≈10^-3^ mutations per genome and generation [97], the half-life of this decay is given by ln 2 /*λ* = ln 2 (2*N_e_*)/(1 + 4*N_e_μ)* ≈ 340-370 generations, well within the time scale of a laboratory evolution experiment. Other simplifications include the assumption that the higher incidence of strongly deleterious mutations in more complex phenotypes holds uniformly across all genotypes with such a phenotype. More sophisticated models are necessary, and they would replace this assumption, for example by allowing for a non-uniform incidence of such mutations. More generally, a statistical sampling theory to estimate quantities such as confidence intervals for estimated information changes also remains an important task for future work.

Despite these caveats, the qualitative considerations I made here suffice to suggest the kind of experimental design needed to estimate phenotypic information changes. Such an experiment would start at the end-point of a previous laboratory evolution experiment, in which a population of organisms or molecules has adapted evolutionarily to a novel environment, such as one containing a specific antibiotic. The experiment would establish two evolving populations. The first derives from a single pre-evolution (ancestral) genotype. It is evolved in the ancestral environment (e.g., without antibiotic). The second population is derived from a single post-evolution (adapted) genotype. It is evolved in the novel environment (e.g., with antibiotic). After allowing both populations to evolve for a sufficient number of generations to approach approximate mutation-selection balance, one would sequence *n* randomly chosen individuals from each population. I showed that for maximal discrimination of information content, it is best if *n* exceeds *N_e_μ* by far in both populations, and that *N_e_μ* itself is greater than one (1≪ *N_e_μ* ≪ *n*). Given limits of sequencing technology, this amounts to conducting the experiment with modest population sizes and mutation rates, but it is entirely feasible. For example, in *E.coli* (μ≈10^-3^), a population of *N_e_*=10^4^ individuals yields *N_e_μ*≈ 10. Current technology allows the sampling conditions to be met, because one can readily sequence *n*>100>>*N_e_μ* complete genomes from microbial populations. The number of different alleles (genomes) sampled from the two populations can then be used to estimate the information difference. An experiment of this kind might be even easier whenever one evolves individual genes rather than whole organisms, because one can then sequence many more alleles, and mutation rates can be manipulated more easily.

Colloquially, evolutionary innovations are viewed as one-of-a-kind, historically singular (and thus rare) qualitative changes, such as *E.coli’s* newfound ability to metabolize citrate after more than 30,000 generations of laboratory evolution [98]. They are distinguished from mere adaptations, such as the gradual increase of a bacterium’s cell division rate in experimental evolution. However, there is no distinction between the two from an information-theoretic perspective. A genotype space that is sufficiently rich, for example that of all proteins of modest length (e.g., 100 amino acids), will harbor a wide variety of different phenotypes, such as proteins that catalyze most chemical reactions important to life. The information content of each is a well-defined number, even though this number may be hard to estimate. Whether a population’s adaptation involves a gradual increase in an enzyme’s activity, or the discovery of an enzyme with a new enzymatic activity, the population merely accesses a new set of genotypes. Innovations may entail larger innovation gains then mere adaptations, but not even that is necessary. For example, in enzymes, where higher catalytic rates require greater phenotypic complexity [75, 95], extremely high rates may be achieved gradually but may require very large information gains.

Because the framework I use is so simple and general, it can speak to diverse issues in evolutionary biology. One of them is the question whether some phenotypes are more evolvable, that is, whether they harbor greater potential to give rise to novel and beneficial phenotypes through Darwinian evolution. A general pattern that emerged from past work is that this potential increases with the number of genotypes that encode a phenotype. Genotypes are usually organized into large genotype networks. The larger such a network is, the greater is the proportion of genotype space that an evolving population can explore, and the greater the population’s chances to originate novel phenotypes phenotypes [67, 71-73]. In information-theoretic terms, low-complexity phenotypes – those encoded by more genotypes – thus harbor greater evolutionary potential. A possible exception can occur when a set of genotypes with the same phenotype is fragmented into multiple subsets (Figure 1d) which cannot be reached from one another by single phenotype-preserving genetic changes [99, 100]. However, in most systems that have been studied, such fragmentation is either absent, creates one dominant genotype network and many smaller networks, or it can be overcome by a combination of large populations, high mutation rates, or strong genetic drift [101-103].

Another, related issue regards the role of DNA or gene duplication in phenotypic innovation. I show that such duplication decreases a phenotype’s informational complexity, because it increases the number of genotypes encoding the phenotype. This holds both in terms of the absolute amount of information and the information content per bit, nucleotide, or any other relevant system part. The latter measure takes into account that duplication changes the size and dimension of a genotype space. And because low phenotypic complexity can come with greater evolutionary potential, the information-theoretic framework can help explain why duplication increases this potential. What is more, it can make quantitative predictions about this potential. For example, the evolutionary benefits of duplication may be greatest for phenotypes with high complexity before duplication, because in such phenotypes, complexity is reduced most strongly after duplication (equation 5).

A third issue is the controversial question whether “progress” occurs during life’s evolution [104]. Progress is not necessarily the same as adaptive evolution. For example adaptive evolution can be regressive [30], as illustrated by many symbiotic and parasitic species that experienced adaptive trait losses. The *Buchnera aphidicola* endosymbionts of pea aphids have dramatically reduced numbers of metabolic enzymes [105]. Threespine stickleback fish have adaptively lost defensive pelvic structures, by losing a regulatory DNA region for the *Pituitary homeobox transcription factor* 1 (Pitx1) [106]. And *E.coli* can lose ribose catabolism during laboratory evolution [107]. The information-theoretic framework allows a simple definition of progress: an increase in phenotype complexity. What is more, the framework can also be used to quantify a loss of such complexity during regressive evolution and trait loss. And the transcription factor binding data illustrates that lowering complexity may not even require degenerative evolution. Consider a gene that acquires regulation by a new transcription factor and loses regulation by an old factor. If the new regulatory phenotype can be realized by more genotypes (transcription factor binding sites), it has lower complexity. We are a long way from quantifying how much more complex an elephant is than *E.coli*, but with information theory, we can at least start to ask quantitative questions about some enduring topics in evolutionary biology, such as evolvability, duplication, and evolutionary progress.

## Methods

### Transcription factor binding data

The experimental data set of 187 mouse transcription factors I analyze here is based on experimental protein binding data from [28] and [29]. It was obtained from the UniPROBE [108](104 transcription factors) and the CIS-BP databases [29] (83 transcription factors) by [88] who identified mouse transcription factors to be included in the data set by several quality criteria, including the requirement that each transcription factor’s DNA binding was assessed on two different kinds of protein binding microarrays, that the factor bound a sufficient number of sequences to permit an analysis of epistasis, and that it bound at least one sequence with an E-score>0.45 [88]. The E-score is related to the Wilcoxon-Mann-Whitney test statistic, and ranges between -0.5 and 0.5 (strongest binding). It is a proxy for a transcription factor’s relative DNA affinity to a binding site [28, 109].For my qualitative analysis, I consider sites bound if E>0.35, because this is a conservative threshold with a small false discovery rate (FDR<0.001) of bound sites [28].

To calculate the information content of a set of DNA sequences of length *L* that bind the same transcription factor under the assumption of additivity across the *L* sites, I first computed the frequencies of all four nucleotides 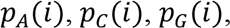 and *p_T_(i)* for each of the *L* positions(1 ≤ *i ≤ L)*. I then computed the entropy for each site as 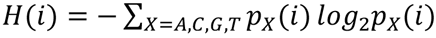, which has a maximum of two if all nucleotides are equally likely to occur at position *i*, and its minimal value of zero if only one nucleotide occurs. The total entropy of a binding site is then simply the sum 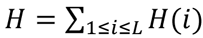, and the total information content is given by *2L — H*, because each of the *L* positions contains maximally two bits. This analysis tacitly assumes that all nucleotides occur at equal frequency, i.e., that no DNA composition bias exists in the underlying genotype space.

### Estimating information gains or changes from sequence data

To estimate *θ_i_* (where *i= ‘New’* or *i=’Old’*, corresponding to populations with the old and new phenotype), the number of different alleles *k_i_*observed in a population sample of *n* sequences is an important quantity. It can be viewed as one realization of a random variable *K_i_* with probability distribution [92, 110]

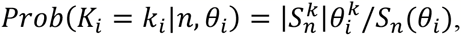

where 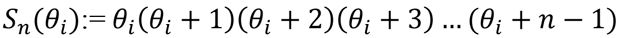 and 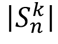 is the absolute value of a Sterling number of the first kind [111], which can be computed numerically. From this distribution, one can compute the expected number of different alleles *F(K_i_)* as

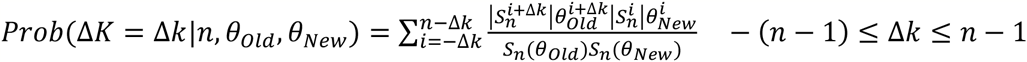

It turns out that for any value of *n* and *k_i_* observed in a sequencing experiment, solving the above equation for *θ_i_* (which can be done numerically) yields not only a maximum likelihood estimator 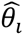 for *θ_i_*, but 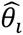 is also a sufficient estimator, that is, one cannot improve it by considering additional information, such as the frequency of each allele in the sample [92 9.5]. In addition, because the distribution of *K_i_* itself is known, one can also compute the distribution of the difference 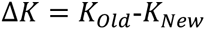 as the following convolution

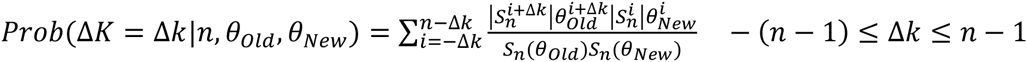

This distribution is valuable, because it helps identify the minimal difference in information content between two phenotypes that can be resolved from a given sequencing experiment. For example, it allows us to test the null-hypothesis that phenotype *P_New_* harbors more information than *P_Old_*, such that one would expect to observe more allelic variants *k_1_* than *k_2_*, and thus that 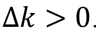 If the null hypothesis is correct, it is possible that one observes by chance alone a value of 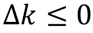 i.e., fewer alleles in population 2, which happens with probability 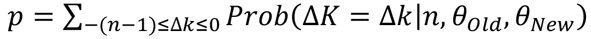. This is the type I error of the hypothesis test, or the probability to falsely reject the null hypothesis. The data in Figure 3 is based on this probability.

## Acknowledgments

I would like to thank Joshua Payne, José Aguilar Rodrigues, and Rzgar Hosseini for valuable discussions on the data analyzed here. I also acknowledge support by Swiss National Science Foundation grant 31003A_146137, by an EpiphysX RTD grant from SystemsX.ch, as well as by the University Priority Research Program in Evolutionary Biology at the University of Zurich.

## References

1 Tkacik, G. and Bialek, W. (2016) Information Processing in Living Systems. In Annual Review of Condensed Matter Physics, Vol 7 (Marchetti, M.C. and Sachdev, S., eds), pp. 89–117

2 Smith, J.M. (2000) The concept of information in biology. Philosophy of Science 67, 177–194

3 Frank, S.A. (2012) Natural selection. V. How to read the fundamental equations of evolutionary change in terms of information theory. Journal of Evolutionary Biology 25, 2377–2396

4 Griffiths, P.E. (2001) Genetic information: A metaphor in search of a theory. Philosophy of Science 68, 394–412

5 van Baalen, M. (2013) Biological information: why we need a good measure and the challenges ahead. Interface Focus 3

6 Taylor, S.F., et al. (2007) Information and fitness. arXiv preprint arXiv:0712.4382

7 Wallace, R. and Wallace, R.G. (1998) Information theory, scaling laws and the thermodynamics of evolution. Journal of Theoretical Biology 192, 545–559

8 Kimura, M. (1961) Natural selection as a process of accumulating genetic information in adaptive evolution. Genetical Research 2, 127-&

9 Haldane, J.B.S. (1957) The cost of natural selection. Genetics 55, 511–524

10 Shannon, C. (1948) A mathematical theory of communication. Bell Systems Technical Journal 27, 379–423

11 Cohen, D. (1966) Optimizing reproduction in a randomly varying environment. Journal of Theoretical Biology 12, 119-&

12 Hoffmann, R.J. (1978) Environmental uncertainty and evolution of physiological adaptation in Colias butterflies. American Naturalist 112, 999–1015

13 Donaldson-Matasci, M.C., et al. (2010) The fitness value of information. Oikos 119, 219–230

14 Donaldson-Matasci, M.C., et al. (2013) When Unreliable Cues Are Good Enough. American Naturalist 182, 313–327

15 McNamara, J.M. and Dall, S.R.X. (2010) Information is a fitness enhancing resource. Oikos 119, 231–236

16 Bergstrom, C.T., et al. (2004) Shannon information and biological fitness.

17 Lachmann, M. and Bergstrom, C.T. (2004) The disadvantage of combinatorial communication. Proceedings of the Royal Society B-Biological Sciences 271, 2337–2343

18 Rivoire, O. and Leibler, S. (2011) The value of information for populations in varying environments. Journal of Statistical Physics 142, 1124–1166

19 Tkacik, G., et al. (2008) Information capacity of genetic regulatory elements. Physical Review E 78

20 Tkacik, G. and Walczak, A.M. (2011) Information transmission in genetic regulatory networks: a review. Journal of Physics-Condensed Matter 23

21 Wagner, A. (2007) From bit to it: The transformation of information into living matter by metabolic networks. BMC Systems Biology 1, 33

22 de Vladar, H.P. and Barton, N.H. (2011) The statistical mechanics of a polygenic character under stabilizing selection, mutation and drift. Journal of the Royal Society Interface 8, 720–739

23 de Vladar, H.P. and Barton, N.H. (2011) The contribution of statistical physics to evolutionary biology. Trends in Ecology & Evolution 26, 424–432

24 Frieden, B.R., et al. (2001) Population genetics from an information perspective. Journal of Theoretical Biology 208, 49–64

25 Weinberger, E.D. (2002) A theory of pragmatic information and its application to the quasi-species model of biological evolution. Biosystems 66, 105–119

26 Demetrius, L. (1974) Demographic parameters and natural selection. Proceedings of the National Academy of Sciences 71, 4645–4647

27 Iwasa, Y. (1988) Free fitness that always increases in evolution. Journal of Theoretical Biology 135, 265–281

28 Badis, G., et al. (2009) Diversity and complexity in DNA recognition by transcription factors. Science 324, 1720–1723

29 Weirauch, M.T., et al. (2014) Determination and inference of eukaryotic transcription factor sequence specificity. Cell 158, 1431–1443

30 Muller, G.B. and Wagner, G.P. (1991) Novelty in evolution: restructuring the concept. Annual Review of Ecology and Systematics, 229–256

31 Dantas, G., et al. (2008) Bacteria subsisting on antibiotics. Science 320, 100–103

32 Toll-Riera, M., et al. (2016) The genomic basis of evolutionary innovation in Pseudomonas aeruginosa. PLoS Genetics 12, e1006005

33 Wilson, D.S. and Szostak, J.W. (1999) In vitro selection of functional nucleic acids. Annual Review of Biochemistry 68, 611–647

34 Keefe, A.D. and Szostak, J.W. (2001) Functional proteins from a random-sequence library. Nature 410, 715–718

35 Hayden, E., et al. (2015) Intramolecular phenotypic capacitance in a modular RNA molecule. Proceedings of the National Academy of Sciences of the U.S.A., 12444–12449

36 Curtis, E.A. and Bartel, D.P. (2013) Synthetic shuffling and in vitro selection reveal the rugged adaptive fitness landscape of a kinase ribozyme. RNA 19, 1116–1128

37 Jiménez, J.I., et al. (2013) Comprehensive experimental fitness landscape and evolutionary network for small RNA. Proceedings of the National Academy of Sciences of the U.S.A. 110, 14984–14989

38 Mukherjee, S., et al. (2004) Rapid analysis of the DNA-binding specificities of transcription factors with DNA microarrays. Nature Genetics 36, 1331–1339

39 Currin, A., et al. (2015) Synthetic biology for the directed evolution of protein biocatalysts: navigating sequence space intelligently. Chemical Society Reviews 44, 1172–1239

40 Beaudry, A.A. and Joyce, G.F. (1992) Directed evolution of an RNA enzyme. Science 257, 635–641

41 Khersonsky, O., et al. (2009) Directed evolution of serum paraoxonase PON3 by family shuffling and ancestor/consensus mutagenesis, and its biochemical characterization. Biochemistry 48, 6644–6654

42 Kiss, C., et al. (2009) Directed evolution of an extremely stable fluorescent protein. Protein Engineering Design & Selection 22, 313–323

43 Romero, P.A. and Arnold, F.H. (2009) Exploring protein fitness landscapes by directed evolution. Nature Reviews Molecular Cell Biology 10, 866–876

44 Hayden, E.J., et al. (2011) Cryptic genetic variation promotes rapid evolutionary adaptation in an RNA enzyme. Nature 474, 92–95

45 Bratulic, S., et al. (2015) Mistranslation drives the evolution of robustness in TEM-1 beta-lactamase. Proceedings of the National Academy of Sciences of the U.S.A. 112, 12758–12763

46 Salverda, M.L.M., et al. (2011) Initial mutations direct alternative pathways of protein evolution. PLoS Genetics 7, e1001321

47 Szendro, I.G., et al. (2013) Quantitative analyses of empirical fitness landscapes. Journal of Statistical Mechanics-Theory and Experiment, P01005

48 Joyce, G.F. (2004) Directed evolution of nucleic acid enzymes. Annual Review of Biochemistry 73, 791–836

49 Dhar, R., et al. (2014) Increased gene dosage plays a predominant role in the initial stages of evolution of duplicate TEM-1 beta-lactamase genes. Evolution 68, 1775–1791

50 Buckling, A., et al. (2009) The Beagle in a bottle. Nature 457, 824–829

51 Lenski, R.E., et al. (1991) Long-term experimental evolution in E. coli.1. adaptation and divergence during 2,000 generations of evolution. American Naturalist 138, 1315–1341

52 Dhar, R., et al. (2013) Yeast adapts to a changing stressful environment by evolving cross-protection and anticipatory gene regulation. Mol. Biol. Evol. 30, 573–588

53 Zeyl, C. (2005) The number of mutations selected during adaptation in a laboratory population of Saccharomyces cerevisiae. Genetics 169, 1825–1831

54 Fong, S.S., et al. (2005) Parallel adaptive evolution cultures of Escherichia coli lead to convergent growth phenotypes with different gene expression states. Genome Research 15, 1365–1372

55 Fares, M.A., et al. (2002) Endosymbiotic bacteria - GroEL buffers against deleterious mutations. Nature 417, 398–398

56 Montville, R., et al. (2005) Evolution of mutational robustness in an RNA virus. PLoS Biology 3, 1939–1945

57 Reboud, X. and Bell, G. (1997) Experimental evolution in Chlamydomonas .3. Evolution of specialist and generalist types in environments that vary in space and time. Heredity 78, 507–514

58 Rifkin, S.A., et al. (2005) A mutation accumulation assay reveals a broad capacity for rapid evolution of gene expression. Nature 438, 220–223

59 Denver, D.R., et al. (2000) High direct estimate of the mutation rate in the mitochondrial genome of Caenorhabditis elegans. Science 289, 2342–2344

60 Maynard Smith, J. (1970) Natural selection and the concept of a protein space. Nature 222, 563 – 564

61 Roscoe, B.P., et al. (2013) Analyses of the effects of all ubiquitin point mutants on yeast growth rate. Journal of Molecular Biology 425, 1363–1377

62 Huang, Z. and Szostak, J.W. (2003) Evolution of aptamers with secondary structures from a new specificity and new secondary structures from an ATP aptamer. RNA-A Publication of the RNA Society 9, 1456–1463

63 Araya, C.L., et al. (2012) A fundamental protein property, thermodynamic stability, revealed solely from large-scale measurements of protein function. Proceedings of the National Academy of Sciences of the United States of America 109, 16858–16863

64 Schuster, P., et al. (1994) From sequences to shapes and back - a case-study in RNA secondary structures. Proceedings of the Royal Society of London Series B 255, 279–284

65 Lipman, D. and Wilbur, W. (1991) Modeling neutral and selective evolution of protein folding. Proceedings of the Royal Society of London Series B 245, 7–11

66 Rodrigues, J.F.M. and Wagner, A. (2009) Evolutionary plasticity and innovations in complex metabolic reaction networks. PLoS Computational Biology 5

67 Payne, J.L. and Wagner, A. (2014) The robustness and evolvability of transcription factor binding sites. Science 343, 875–877

68 Hosseini, S.-R., et al. (2015) Exhaustive analysis of a genotype space comprising 10^15^ central carbon metabolisms reveals an organization conducive to metabolic innovation. PLoS Comput Biol 11, e1004329

69 Manrubia, S. and Cuesta, J.A. (2015) Evolution on neutral networks accelerates the ticking rate of the molecular clock. Journal of the Royal Society Interface 12, 20141010

70 Ancel, L.W. and Fontana, W. (2000) Plasticity, evolvability, and modularity in RNA. Journal of Experimental Zoology/Molecular Development and Evolution 288, 242–283

71 Wagner, A. (2008) Robustness and evolvability: a paradox resolved. Proceedings of the Royal Society of London Series B-Biological Sciences. 275, 91–100

72 Ferrada, E. and Wagner, A. (2008) Protein robustness promotes evolutionary innovations on large evolutionary time scales. Proceedings of the Royal Society of London Series B-Biological Sciences. 275, 1595–1602

73 Greenbury, S.F., et al. (2014) A tractable genotype–phenotype map modelling the self-assembly of protein quaternary structure. Journal of the Royal Society Interface 11, 20140249

74 Wootton, J.C. (1994) Non-globular domains in protein sequences: automated segmentation using complexity measures. Computers & Chemistry 18, 269–285

75 Szostak, J.W. (2003) Molecular messages. Nature 423, 689–689

76 Cover, T.M. and Thomas, J.A. (2006) Elements of information theory. Hoboken, New Jersey

77 Neylon, C. (2004) Chemical and biochemical strategies for the randomization of protein encoding DNA sequences: library construction methods for directed evolution. Nucleic Acids Research 32, 1448–1459

78 Hartl, D.L. and Clark, A.G. (2007) Principles of population genetics. Sinauer Associates

79 Ohno, S. (1970) Evolution by gene duplication. Springer

80 Conant, G.C. and Wolfe, K.H. (2008) Turning a hobby into a job: How duplicated genes find new functions. Nature Reviews Genetics 9, 938–950

81 Innan, H. and Kondrashov, F. (2010) The evolution of gene duplications: classifying and distinguishing between models. Nature Reviews Genetics 11, 97–108

82 Theissen, G. (2001) Development of floral organ identity: stories from the MADS house. Current Opinion in Plant Biology 4, 75–85

83 Wagner, A. (2008) Gene duplications, robustness and evolutionary innovations. Bioessays 30, 367–373

84 Wray, G.A. (2007) The evolutionary significance of cis-regulatory mutations. Nature Reviews Genetics 8, 206–216

85 Tuch, B.B., et al. (2008) The evolution of combinatorial gene regulation in fungi. PLoS Biology 6, 352–364

86 Prud&homme, B., et al. (2007) Emerging principles of regulatory evolution. Proceedings of the National Academy of Sciences of the United States of America 104, 8605–8612

87 Carroll, S.B., et al. (2001) From DNA to diversity. Molecular genetics and the evolution of animal design. Blackwell

88 Aguilar-Rodriguez, J.P., J.A. and Wagner, A. (2016) 1000 empirical adaptive landscapes and their navigability. (submitted).

89 Weaver, D.C., et al. (1999) Modeling regulatory networks with weight matrices. Pacific Symposium on Biocomputing 4, 112–123

90 Wagner, A. (2011) The molecular origins of evolutionary innovations. Trends in Genetics 27, 397–410

91 Kimura, M. (1983) The neutral theory of molecular evolution. Cambridge University Press

92 Ewens, W.J. (2012) Mathematical Population Genetics 1: Theoretical Introduction. Springer Science & Business Media

93 Adami, C., et al. (2000) Evolution of biological complexity. Proceedings of the National Academy of Sciences 97, 4463–4468

94 Strelioff, C.C., et al. (2010) Evolutionary dynamics, epistatic interactions, and biological information. Journal of Theoretical Biology 266, 584–594

95 Carothers, J.M., et al. (2004) Informational complexity and functional activity of RNA structures. Journal of the American Chemical Society 126, 5130–5137

96 Fontana, W. and Schuster, P. (1998) Continuity in evolution: On the nature of transitions. Science 280, 1451–1455

97 Lee, H., et al. (2012) Rate and molecular spectrum of spontaneous mutations in the bacterium Escherichia coli as determined by whole-genome sequencing. Proceedings of the National Academy of Sciences of the United States of America 109, E2774–E2783

98 Blount, Z.D., et al. (2008) Historical contingency and the evolution of a key innovation in an experimental population of Escherichia coli. Proceedings of the National Academy of Sciences 105, 7899–7906

99 Greenbury, S.F., et al. (2014) A tractable genotype-phenotype map modelling the self-assembly of protein quaternary structure. Journal of the Royal Society Interface 11

100 Schaper, S., et al. (2012) Epistasis can lead to fragmented neutral spaces and contingency in evolution. Proceedings of the Royal Society B-Biological Sciences 279, 1777–1783

101 Weinreich, D.M. and Chao, L. (2005) Rapid evolutionary escape by large populations from local fitness peaks is likely in nature. Evolution 59, 1175–1182

102 Weissman, D.B., et al. (2009) The rate at which asexual populations cross fitness valleys. Theoretical Population Biology 75, 286–300

103 Wagner, A. (2011) The origins of evolutionary innovations. A theory of transformative change in living systems. Oxford University Press

104 Ruse, M. (1993) Evolution and Progress. Trends in Ecology & Evolution 8, 55–59

105 Thomas, G.H., et al. (2009) A fragile metabolic network adapted for cooperation in the symbiotic bacterium Buchnera aphidicola. BMC Systems Biology 3

106 Chan, Y.F., et al. (2010) Adaptive evolution of pelvic reduction in sticklebacks by recurrent deletion of a Pitx1 enhancer. Science 327, 302–305

107 Cooper, V.S., et al. (2001) Mechanisms causing rapid and parallel losses of ribose catabolism in evolving populations of Escherichia coli. Journal of Bacteriology 183, 2834–2841

108 Newburger, D.E. and Bulyk, M.L. (2009) UniPROBE: an online database of protein binding microarray data on protein–DNA interactions. Nucleic Acids Research 37, D77–D82

109 Berger, M.F., et al. (2006) Compact, universal DNA microarrays to comprehensively determine transcription-factor binding site specificities. Nature Biotechnology 24, 1429–1435

110 Ewens, W.J. (1972) The sampling theory of selectively neutral alleles. Theoretical population biology 3, 87–112

111 Abramowitz, M. and Stegun, I. (1972) Handbook of mathematical functions. Dover

